# A mechanical modelling framework to study endothelial permeability

**DOI:** 10.1101/2023.07.28.551049

**Authors:** Pradeep Keshavanarayana, Fabian Spill

## Abstract

The inner lining of blood vessels, the endothelium, is made up of endothelial cells. Vascular endothelial (VE)-cadherin protein forms a bond with VE-cadherin from neighbouring cells (homophilic bond) to determine the size of gaps between the cells and thereby regulate the size of particles that can cross the endothelium. Chemical cues such as Thrombin, along with mechanical properties of the cell and extracellular matrix (ECM) are known to affect the permeability of endothelial cells. Abnormal permeability is found in patients suffering from diseases including cardiovascular diseases, cancer, and COVID-19. Even though some of the regulatory mechanisms affecting endothelial permeability are well studied, details of how several mechanical and chemical stimuli acting simultaneously affect endothelial permeability are not yet understood.

In this article, we present a continuum-level mechanical modelling framework to study the highly dynamic nature of the VE-cadherin bonds. Taking inspiration from the catch-slip behaviour that VE-cadherin complexes are known to exhibit, we model VE-cadherin homophilic bond as cohesive contact with damage following a traction-separation law. We explicitly model the actin-cytoskeleton, and substrate to study their role in permeability. Our studies show that mechano-chemical coupling is necessary to simulate the influence of the mechanical properties of the substrate on permeability. Simulations show that shear between cells is responsible for the variation in permeability between bi-cellular and tri-cellular junctions, explaining the phenotypic differences observed in experiments. An increase in the magnitude of traction force that endothelial cells experience results in increased permeability, and it is found that the effect is higher on stiffer ECM. Finally, we show that the cylindrical monolayer exhibits higher permeability than the planar monolayer under unconstrained cases. Thus, we present a contact mechanics-based mechano-chemical model to investigate the variation in permeability of endothelial monolayer due to multiple loads acting simultaneously.

## 1 Introduction

The vascular network is responsible for the transport of blood, and other essential nutrients across the body via arteries, veins and capillaries. Arteries carry the blood away from the heart, veins bring the blood back to the heart while capillaries act as a bridge between arteries, and veins and ensure the delivery of blood, nutrients, and oxygen to the surrounding tissue. Capillaries are part of the micro-circulation and are the smallest blood vessels in the human body. The blood vessels are made up of a specialised type of cells called vascular endothelial (VE) cells. They form a network of cylindrical monolayers, termed endothelium, allowing fluid to flow within. In the case of arteries and veins, the endothelium is surrounded by smooth muscle tissues while capillaries are made up of a single layer of endothelial cells, which is the focus of this article and study.

In a healthy vascular system, the homeostasis of capillaries is implied by a well-regulated bi-directional transport of material between endothelium and the surrounding tissues. The transport can occur via two pathways: transcellular (through the cell), and paracellular (between the cells) ^1^. Transcellular transport involves endocytosis (into the cell), and exocytosis (out of the cell). These processes are highly selective and it is found that only a few micronutrients such as Vitamin B12 follow the transcellular pathway ^2^. On the other hand, the paracellular pathway is regulated by several intercellular junctional proteins that regulate the gap size between the cells. Materials passing the endothelium via the paracellular pathway have to overcome the barrier properties of these inter-junctional proteins ^1^. One of the intercellular junctions is termed the adherens junction and is commonly found in the vascular endothelium. VE-cadherin proteins are the major component of adherens junctions. They form a homophilic bond with VE-cadherin proteins on the neighbouring cells and dictate the size of material that can pass the endothelium ^3^. It is to be noted that the VE-cadherin-VE-cadherin homophilic bond is highly dynamic in nature, resulting in the constant opening and closing of gaps due to changes in cellular and extracellular properties. The number and size of gaps present decide the amount of material that can successfully cross the endothelium, which is commonly termed (vascular) permeability. Thus, an increase in the amount of material that can overcome the endothelial barrier indicates higher endothelial permeability.

Dysregulation of VE-cadherin homophilic bonds may lead to either larger gaps or changes in the frequency of gap formation or gaps not being formed, leading to abnormal permeability. Some cases of diseased states where abnormal permeability is found are cancer ^4^, atherosclerosis ^5^, asthma ^6^, and COVID-19^7^. Cancer metastasis, a key characteristic of cancer progression, involves the spread of cancer cells from their origin to other body parts via the vascular network. This metastasis is facilitated by abnormal vascular permeability, which allows for the intravasation (entry into blood vessels) and extravasation (exit from blood vessels) of cancer cells. Similarly, abnormal permeability plays a significant role in atherosclerosis, a prevalent cardiovascular disease in humans. This condition is characterised by the infiltration of fat and cholesterol molecules from the blood through the endothelial layer, subsequently accumulating on the surrounding membrane. This accumulation narrows the blood vessels, disrupting normal blood flow, and in severe cases, may result in cardiac arrest. Vascular permeability has a major role in both acute and chronic inflammation since leukocytes have to cross the endothelial barrier and reach the tissues. VE-cadherin-based cell-cell junctions are found to play an important role in angiogenesis and vascular morphogenesis ^8, 9^. Thus, it becomes prudent to understand the physiology of variation of permeability of the endothelium. A better knowledge of how VE-cadherin bonds could be regulated helps us to understand the disease pathology and develop cures.

VE-cadherins are downstream connected to the actin-cytoskeleton via the catenin and vinculin family of proteins ^10^, as shown in Fig. 1. Thus, regulation of the actin-cytoskeleton is expected to affect the binding properties of the VE-cadherins as well. Experiments have shown how modifying the contractility of actin-myosin stress fibres via Rho/ROCK pathways affects permeability ^11^. Some chemical cues such as thrombin increase permeability ^12^ while S1P reduces permeability and stabilises the cell-cell junctions ^13^. In addition to chemical cues, mechanical cues are also found to affect cell-cell junctions. Properties of the micro-environment such as stiffness of the substrate ^14^, geometric properties of the substrate ^15^, and mechanical stimuli such as cyclic stretch ^16^ are all known to affect permeability. But *in-vivo*, chemical and mechanical stimuli act simultaneously and there is a coupling between them. Experiments carried out *in-vitro* usually study either the effect of chemical or mechanical stimuli individually. In this regard, mathematical models that can integrate different features play an important role in bridging the understanding between individual responses observed experimentally.

**Figure 1:**
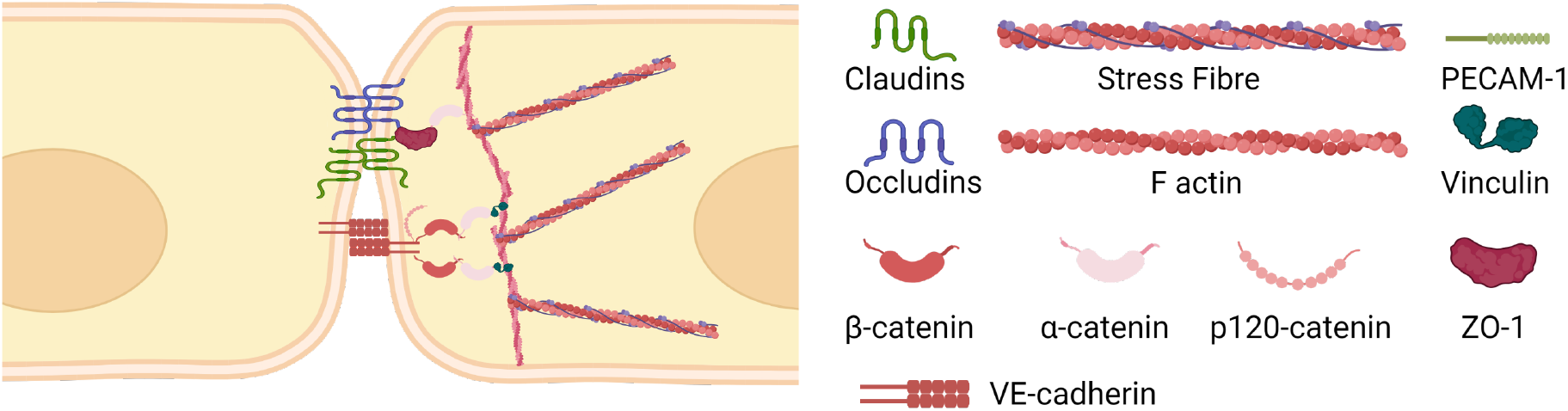
Schematic of proteins involved in cell-cell junction mechanotransduction. VE-cadherins from neighbouring cells form a homophilic bond as part of the adherens junctions. VE-cadherins are attached to the Catenin family of proteins, which under tension, are connected to the cytoskeleton via Vinculin. Claudins and Occludins present in between the cells form tight junctions, that are mainly found in the Brain vascular network. Created with biorender.com

Over the past few decades, several mathematical models have been developed to understand the behaviour of endothelial cells in differing mechanical and chemical environments. Many models focused on simulating angiogenesis either from mechanical or mechano-chemical perspectives ^17–19^. In order to study the behaviour of monolayers as opposed to individual cells, the 2D cellular Potts model was used along with a positive feedback signal between cellular movement and polarity to simulate flow fields for differing cell densities ^20^. A dynamic model of cell steering showed the effect of low-level gene changes on the high-level migration of a monolayer ^21^. The effect of heterogeneity of substrate properties on the strain profiles of the endothelial monolayer was studied by developing a linear elastic model of the nucleus, cytoplasm, matrix and VE-cadherin ^22^.

In recent times, there has been a renewed interest in developing mathematical models to understand the physiology of gap dynamics. The agent-based model ^23^ was one of the first models developed to study the effect of mechanical properties of cells to study the dynamics of endothelial permeability. Considering actin stress fibres as visco-elastic springs and adhesion dynamics modelled to follow catch-slip bond law, the model was able to predict the dynamics of cell-cell junctions. A discrete continuum hybrid model ^24^ was developed to study the effect of calcium waves on cell-cell junctions. It was noted that the gaps were higher at the vertices of cells and thus cancer cells extravasate at these locations over edges. Recently, a continuum model was developed to simulate the three-way feedback between VE-cadherins, RhoA contractility and Rac-1 derived actin polymerisation ^25^. The myosin-dependent force generation was balanced by the force due to VE-cadherins in addition to being coupled to signalling and cell stress. However, the models developed so far have not considered the effect of several types of mechanical stimuli acting simultaneously, such as traction due to the flow of blood, cell contractility, and hemodynamic pressure, on the endothelium. The effect of mechanical properties of VE-cadherin and the role of cell-substrate junctions in regulating endothelial permeability have not been studied either. In this regard, we have developed a novel continuum-level contact-mechanics-based mathematical model that couples VE-cadherin to Ca^2+^ based stress fibre contractility and cell-substrate adhesion. VE-cadherins are thus mechanical entities that can form and break contact with VE-cadherins from neighbouring cells following the principle of traction-separation law. We have been able to study the effect of different loads acting simultaneously and the role of substrate stiffness on permeability using this model. In the following sections, we will introduce the model and show several numerical examples that demonstrate how different mechanical and chemical stimuli affect permeability.

## 2 Mathematical model

In this article, we introduce a novel mathematical model where the cell-cell contact force is balanced by the actin-cytoskeleton contractile force and cell-substrate contact force, as shown in Fig. 2a. In the current continuum modelling framework, the VE-cadherin bonds are formulated as contact cohesive surfaces. This allows us to simulate the zip-like behaviour observed in cell-cell junctions ^26^ and also be able to extend the geometry to 3D, unlike earlier models that focused mainly on 2D geometries. Bonds between the VE-cadherins of neighbouring cells are modelled to be formed when two cell surfaces come in contact with each other and broken when they lose contact, determined by the separation distance between them.

**Figure 2:**
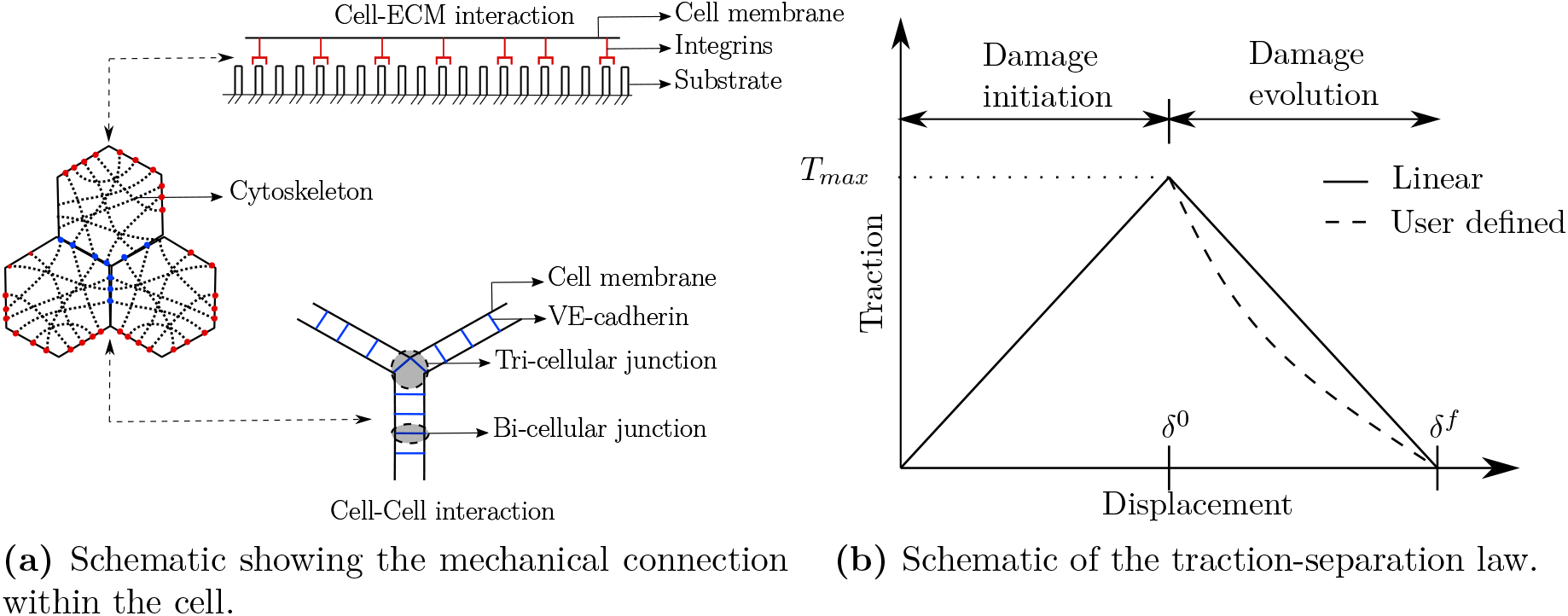
a) Cytoskeleton connects integrins on one end and VE-cadherins on the other. b) Traction separation law used as a mechanical equivalent of catch-slip bond. Damage initiation is assumed to be linear and evolution can be either linear or any other user-defined function. In this article, damage evolution also is assumed to be linear. *δ*^0^ indicates the relative displacement at which the damage is initiated in each direction, and *δ^f^*, the effective relative displacement at complete damage.

The catch-slip bond law that has been used in literature for simulating cell-cell adhesion dynamics considers a linear combination of catch and slip zones exhibited by the cell-cell adhesion ^23^. Simulations show that while stretching the cell-cell junctions, the rate of binding increases up to a particular limit, and then reduces. Recent *in-vitro* experiments performed on a single cell-cell junction have shown that the force-displacement curve exhibits a linear rise up to a point and then drops after reaching a peak ^27^. In this regard, we hypothesise traction-separation law as a mechanical equivalent of the catch-slip bond law, explained in detail in section 2.1 and use this on top of cohesive surfaces to dictate how surfaces should damage (disassembly of bonds). The model has been implemented in ABAQUS, which is a commercial finite element solver ^28^. Files with geometry, material definitions and other algorithmic implementations (inp, UMAT, UAMP) are distributed as open-source and can be downloaded via the link provided in the supplemental material section S1.

### 2.1 VE-cadherin model

The traction-separation law for the damage of cell-cell junction could be split into two zones, one for the damage initiation and the other for the damage evolution, as can be seen in Fig. 2b, which could be compared to the binding and unbinding zones present in a catch-slip bond law. When the surfaces in contact are stretched away up to *δ*^0^, the bonds stabilise with stretch and the traction force increases. When the stretch reaches *δ*^0^, the damage is said to be initiated and the bond starts to destabilise. Further, stretch of the bond at which there is complete damage is indicated by *δ^f^*. A linear constitutive relation between stress and strain in the VE-cadherin bond prior to damage initiation is assumed as given in Eq. (1).

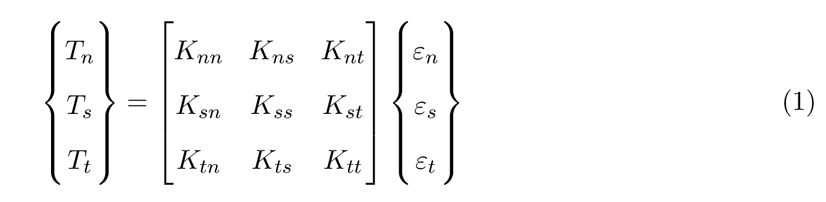

where *T* is the traction, K is the stiffness and *ε* is the strain given in the matrix notation. Subscript *n* indicates the normal direction and *s*, and *t* indicates the shear directions. As can be seen from Fig. 2b damage could be initiated either upon reaching a relative displacement of *δ*^0^ (maximum separation criteria) or a traction of *T_max_* (maximum traction criteria). In this article, we use the maximum separation criteria as given in Eq. (2) for the quantitative analyses,

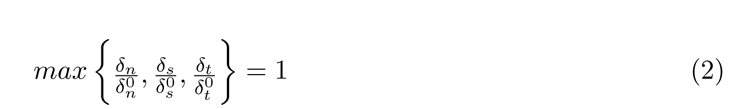

where *δ*^0^_*n*_,*δ*^0^_*n*_,*δ*^0^_*n*_ are the maximum values of separation allowed for damage initiation in normal, and shear directions, and *δ_n_*, *δ_s_*, *δ_t_* are the current separation values in their respective directions, but can equivalently be analysed using maximum traction criteria. Once the damage is initiated, its evolution can follow linear, exponential or any other user-defined response, as can be seen in Fig. 2b. In this article, we use a linear evolution law, where the damage variable D, representing the overall damage in the material (equivalent to slip zone) follows Eq. (3). The initial value of D is 0, which increases to a value of 1 upon complete damage.

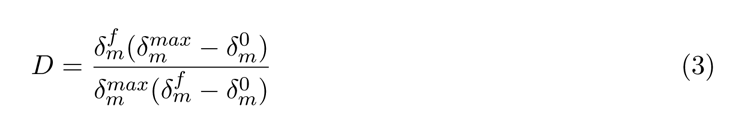

Here, 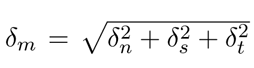 is the effective separation. *δ^f^* is the effective stretch at complete damage, and *δ^max^* indicates the maximum value of effective separation attained during the loading history. The effective displacement at the damage initiation (*δ*^0^), without loss of generality, assuming *δ*^0^ = *δ*^0^, can be evaluated, following ^29^, as

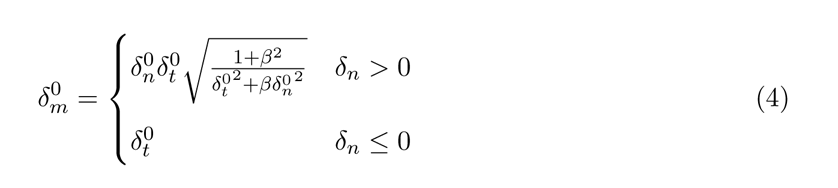

where, 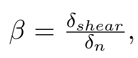, and *δ_shear_* represents the norm of the vector defining the tangential relative displacements (*δ_s_*, *δ_t_*). Readers are referred to ^29^ for further details on derivation and analysis. The stress components (*T_n_*, *T_s_*, *T_t_*) are affected due to damage according to Eq. (5), where (*T*^̄^*_n_*, *T*^̄^*_s_*, *T*^̄^*_t_*) are the stress components predicted by elastic traction-separation law prior to damage initiation.

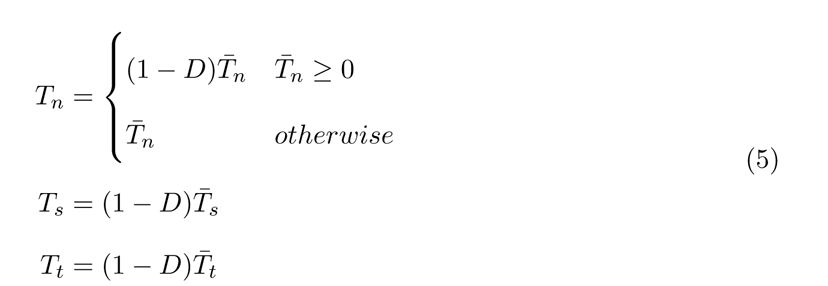

Thus, using Eqs. (1) - (5), VE-cadherin-based cell-cell adhesion has been modelled in this article, and the behaviour is similar to the catch-slip bond observed experimentally. In a catch-slip bond model, by varying the rate of association and dissociation, with force applied on the bond being the independent parameter, a spectrum of bond lifetime vs force curves can be obtained ^23^. Similarly, by changing the values of separation at damage initiation (*δ*^0^) and at complete failure (*δ^f^*), we can obtain a range of traction-separation curves. Even though there is no direct correlation between the lifetime-force curves obtained using the catch-slip bond law and traction-separation curves obtained via contact mechanics-based traction-separation law, physics simulated by both of them are equivalent. More details of parameter variation of traction-separation law are given in the supplemental material section S2. In this article, the value of bond stiffness is chosen such that the maximum value of contact force is in the same range as the bond force observed experimentally ^30^. Additionally, it is to be noted that the lifetime of a bond is non-zero at an initial state where the applied force is zero, while in the case of traction-separation law, the initial condition is chosen to be with a contact force being zero. In addition to cell-cell adhesion, it has been observed that cell-substrate adhesion also behaves as a catch-slip bond ^31^. Hence, to study the role of cell-substrate contact on permeability *in-silico*, we model the integrin-based cell-substrate bond using traction-separation law.

### 2.2 Cytoskeleton model

The VE-cadherins are further connected to the actin-cytoskeleton via the catenin and vinculin family of proteins. In order to include the effect of actin-cytoskeleton on VE-cadherin bonds, we assume the total stress in the cell to comprise two components, active stress (*σ^a^*) and passive stress (*σ^p^*), Eq. (6). Active stress is generated by the stress fibre contractility while passive stress is due to other components in the cytoskeleton.

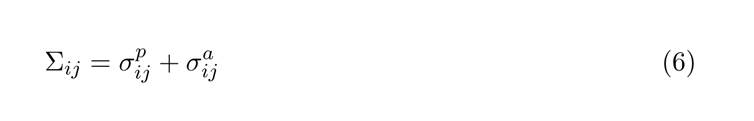

where *i, j* = 1, 2, 3 are indices, following Einstein summation convention. In this article, for simplicity, the passive stress growth is assumed to follow a linear elastic material law, as given in Eq. (7).

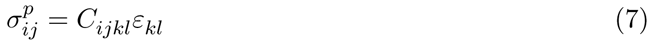

where *C_ijkl_* is the stiffness tensor, which depends on Young’s modulus *E*, and Poisson’s ratio *ν*. The active part of the cytoskeleton is assumed to be composed of stress fibres oriented in discrete angles, and follows a strain-rate dependent growth ^32^, as given in Eq. (8). Readers are referred to ^32–34^ for more information on the active stress growth model used.

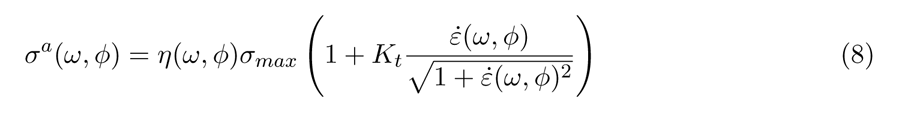

where *η*(*ω, ϕ*) (0 *≤ η*(*ω, ϕ*) *≤* 1) is a scalar representing the stress fibre concentration at a particular orientation (*ω, ϕ*). The angle that the fibre makes with the *x*_3_ axis is given by *ω* and *ϕ* is the angle of the projection of fibre on *x*_1_-*x*_2_ plane with respect to the *x*_1_ axis. Following ^32^, the growth of stress fibre concentration follows an ODE, given in Eq. (9)

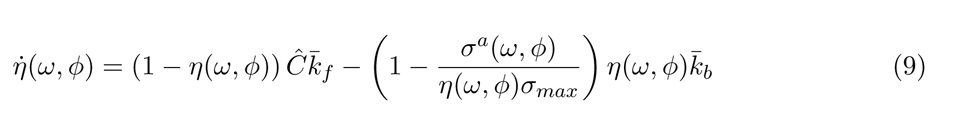

where *k*^̄^*_f_* and *k*^̄^*_b_* are the rate of association and dissociation, and *C*^^^ is the calcium concentration. The maximum stress that the stress fibres can carry is given by *σ_max_*. Upon evaluating the active stress in a given stress fibre oriented at a particular orientation (*ω*,*ϕ*), the active Cauchy stress tensor can be evaluated ^33, 35^ as given in Eq. (10)

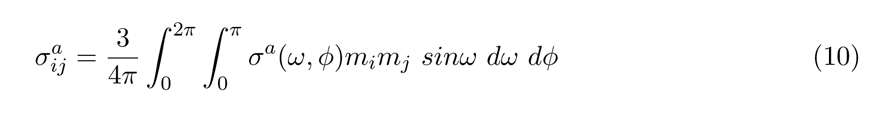

where, *m_i_* and *m_j_* denote the *i*, and *j* components of the generalised unit vector in the direction of the given stress fibre. Further details on derivation are given in the supplemental material section S3. Upon addition of Eqs. (7) and (10), the total Cauchy stress Eq. (6) in the cell can be evaluated. The static mechanical equilibrium equation is then solved using a commercial finite element solver ABAQUS.

### 2.3 Numerical scheme

The finite element method is used to solve the mechanical equilibrium between the cytoskeletal stress and the contact stress generated at the cell-cell and cell-substrate interactions. We use tetrahedral elements, with linear shape functions. Numerical integration of equation Eq. (10) is performed by assuming that there are *N_f_* fibres equally distributed over the cell domain. Numerical integration of the stress fibre growth ODE is performed via the forward Euler method. In the case of small strain formulation, the consistent Jacobian (C) needed for solving the mechanical equilibrium can be evaluated as

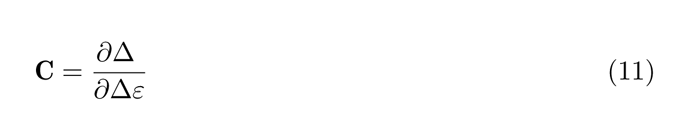

It is to be noted that the Jacobian **C** is not part of the stress update, and is required by the solver only to obtain quadratic convergence of the Newton scheme. Hence, using different methods to evaluate Eq. (11) might only affect the convergence and does not affect the result. As mentioned earlier, in this article, the passive stress growth is assumed to be linear-elastic, and small strain finite element analysis is performed. This could easily be replaced by hyperelastic materials such Neo-Hookean and finite strain analysis can be performed. The procedure to be followed to evaluate Jacobian in the case of finite strain formulation is given in the supplemental material S4.

### 2.4 Permeability of the endothelium

The endothelium is said to be permeable to a particular molecule when it is possible for that molecule to cross the endothelial barrier. As explained earlier, in this article, we consider only the VE-cadherin-regulated paracellular pathway to study permeability. This is an important measure of a core function of the endothelium and a key measure of health and disease. Understanding permeability helps in designing effective drug delivery systems. In experiments, permeability is usually measured in two ways: 1. Using a dye to measure the amount of tracer molecule that has crossed the endothelium via cell-cell junctions ^36^ and 2. By measuring the transendothelial electrical resistance (TEER) ^37^.

Experiments, *in-vitro*, performed on HAECs estimate the permeability of the monolayer by measuring the amount of FITC-avidin tracer that crosses the monolayer via the paracellular gaps ^36^. The sub-endothelial accumulation of the tracer is quantified using a fluorimetric plate reader. The higher amount of tracer collected below the endothelial cell monolayer indicates higher permeability of the monolayer. The tracer molecules are larger than the intracellular pathways, resulting in the quantification of paracellular pathways alone. The size of the tracer molecule used can also indirectly indicate the size of paracellular gaps, which is not possible to be measured using the TEER technique.

Thus, for in *in-silico* experiments, we define permeability (*χ*) as the ratio of the number of open cell-cell junctions (N*_o_*) to the total number of cell-cell junctions (N*_t_*) in the model of a monolayer, Eq. (12). A junction is said to be open if the normal distance between two such surfaces is greater than *δ^f^* or equivalently if the contact force between two surfaces that were in contact earlier becomes 0. The normal distance between two surfaces can imply the gap size, and therefore variation of permeability with respect to gap size can be studied. Cell-Cell junctions in a monolayer can be split into bi-cellular, and tri-cellular junctions where two and three cells are in contact respectively, as shown in Fig. 2a. In the simulations presented in this article, as we perform linear static analysis where all deformation measures are evaluated with respect to the initial configuration, bi-cellular and tri-cellular junctions are also defined with respect to the initial configuration and are not updated with deformation.

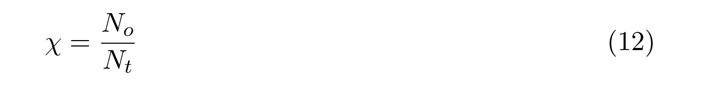

### 2.5 Initial and Boundary conditions

The initial geometry of the cell is considered to be a regular hexagon with sides of 10*µ*m and a thickness of 0.1*µ*m ^38^. The initial condition is assumed to be stress-free, with cells perfectly in contact with the neighbouring cells. The distance between the surfaces in contact is taken to be 0 at the stress-free state. A small-sliding contact formulation with cohesive behaviour for surface-surface contact is defined, with any secondary node experiencing contact taken to be an eligible secondary node for contact. This allows a given master node to form a cohesive bond with any secondary node that comes in contact and satisfies the traction separation law, even though those two nodes may not be in contact at the initial state. Normal contact behaviour is defined as ”hard-contact”, preventing the penetration of cells, and allowing separation of surfaces after contact. The outermost walls of the outermost cells in a given monolayer are assumed to be fixed (all degrees of freedom are constrained). The parameter values chosen are given in the supplemental material section S5. The results presented in this article are for qualitative analysis, and we do not aim to perform quantitative comparisons with any existing literature.

## 3 Results and Discussions

### 3.1 Location of permeability depends on mechanical stimuli

To begin with, we consider a planar monolayer of hexagonal cells. In *in-vitro* experiments performed on planar monolayers, pro-inflammatory agents such as Thrombin are used to induce permeability. The addition of Thrombin activates the RhoA/ROCK pathway causing an increase in contractility which leads to a rise in permeability. In our model, we simulate such behaviour via the Calcium concentration parameter *C*^^^, used to evaluate stress fibre concentration, Eq. (9). Activation of *C*^^^ (non-zero, but uniform over the cell domain), which could be thought of as an increase in the cytoplasmic calcium concentration, results in the growth of active stress (*σ^a^*(*ϕ, ω*)), and cells undergo uniform contraction. This results in a higher strain at the tri-cellular junctions than at bi-cellular junctions. This leads to gaps formed at tri-cellular junctions prior to bi-cellular junctions. But as seen from experiments, cells do not exhibit uniform contraction. Random thermal fluctuations and other non-equilibrium cellular processes occur in the cell ^39^ resulting in a change of shape and its mechanical properties. We hypothesise that this behaviour leads to the random opening and closing of VE-cadherin bonds. In order to simulate such behaviour, we divide the boundary of each cell into a number of smaller divisions and apply random pressure loading on each of these divisions, whose magnitude, we assume, is directly proportional to *C*^^^. Based on the history of the loading on the individual divisions, each of them could be either open or closed independently of the other resulting in a behaviour similar to that observed experimentally. With the application of random pressure loading, we see that strain distribution is now more heterogeneous with local regions of high and low strain present. Based on the load, the strain could be high either at a tri-cellular junction or bi-cellular junction and hence, permeability. In addition to the pressure loading, which is always normal to the surface, random shear loading, parallel to the surface, can also be applied. Strain distribution in case of uniform and random opening of gaps is shown in supplemental material section S7, Figure S3.

Following the definition of permeability as defined in Eq. (12), we can evaluate permeability for the planar monolayer undergoing random opening and closing of VE-cadherin bonds. Due to random loads, cell-cell junctions open and close over time, and reach a steady state as seen in Fig. 3a. Investigating whether the permeability occurs at bi-cellular or tri-cellular junctions, we find that permeability is higher for bi-cellular junctions compared to tri-cellular junctions. Since random pressure loads applied are normal to the surface, the fraction of bi-cellular junctions that break is higher than the tri-cellular junctions resulting in a higher value of *χ* at bi-cellular junctions. Steady-state permeability can be defined by evaluating the average of the last 100 data points (iterations) individually for bi-cellular, and tri-cellular junctions and also for the mean presented as mean*±*SD in Fig. 3b.

**Figure 3:**
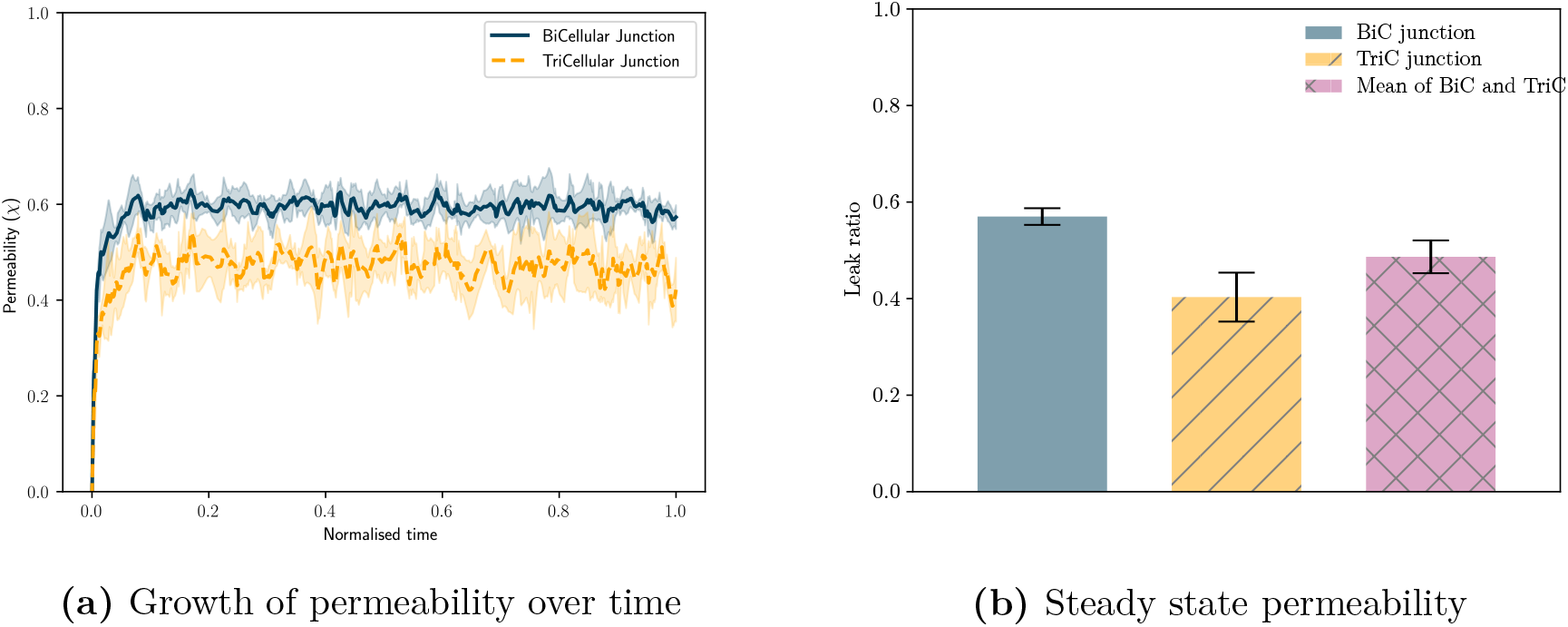
Permeability is defined by the ratio of the number of open junctions to the total number of junctions. (a) Permeability varies randomly over time indicating that the number of open junctions fluctuates over time. (b) Steady-state values of permeability can be evaluated for bi-and tri-cellular junctions and the mean. Error bars indicate mean*±*SD to highlight variability.

Further, we study the variation of permeability by changing the threshold of the normal distance between cells that defines when a gap is considered open. We hypothesise that this is equivalent to changing the size of tracer molecules in an *in-vitro* experiment. Our simulations show that as the criteria for a gap to be considered open is increased, permeability reduces. It is found that this drop is non-linear, as can be seen in Fig. 4a. This shows that the permeability of endothelial monolayer to a 1*µ*m particle is almost 10 times lower than that of a 0.15*µ*m particle and 4 times that of a 0.4*µ*m particle. Thus, with this simulation, based on the size of the leukocyte (*in-vivo*) or tracer molecule (*in-vitro*), we can estimate the chance of crossing the endothelial barrier via the paracellular pathway.

**Figure 4:**
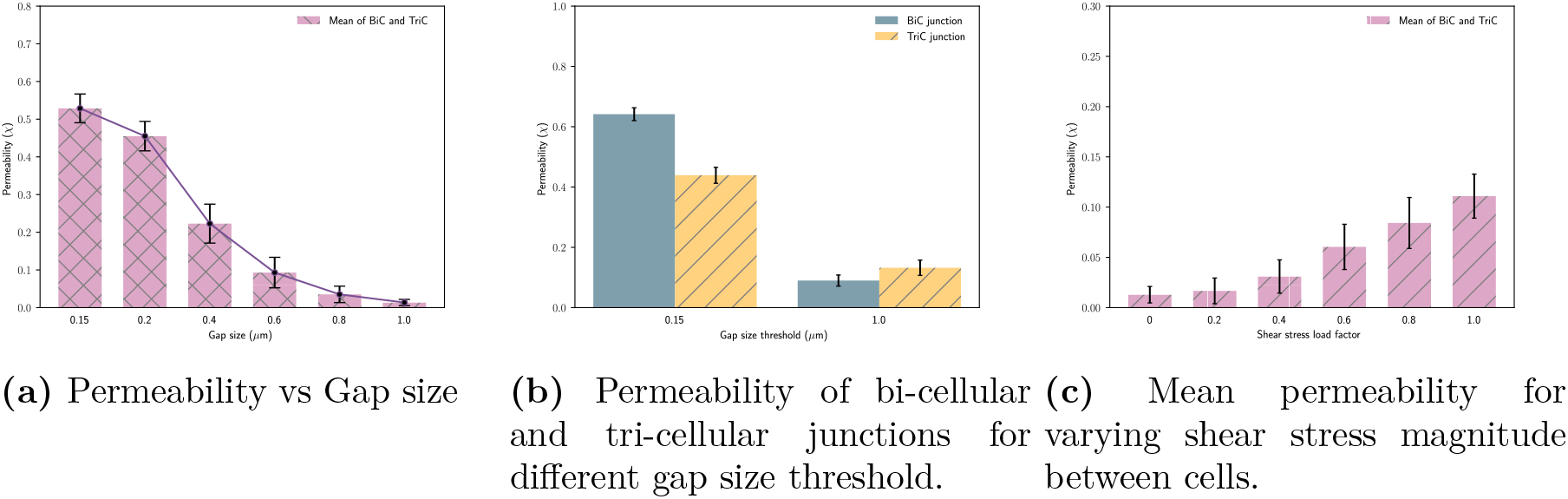
Effect of mechanical stimuli and total damage parameter *δ^f^* on permeability. a) As the gap size threshold increases, permeability reduces exponentially. b) The permeability of a tri-cellular junction is found to be higher than a bi-cellular junction due to shear, only when the gap size threshold is high. c) As the magnitude of shear between cells is increased, the permeability also increases. Gap size threshold considered = 1*µ*m.

*In-vitro* experimental studies have shown that the difference in permeability between bi-and tri-cellular junctions depends on the type of cell, passage number, and tracer molecule used. It was observed that while studying permeability with human aortic endothelial cells (HAECs), bi-cellular junctions showed less involvement compared to that with porcine aortic endothelial cells, and found that during experiments with HAECs a majority of tracer molecules accumulated below tri-cellular junctions ^36^. We, therefore, hypothesise that differences observed due to changes in cell types are due to variations in mechanical stimuli that the cells experience. Hence, we apply shear stress at cell-cell junctions with randomly varying magnitudes in addition to random pressure loading. It was observed that even with shear between cells, for a gap size threshold of 0.15*µ*m, permeability at bi-cellular junctions was higher than at tri-cellular junctions. But, upon increasing the gap size threshold to 1.0*µ*m, we see that the permeability is higher at tri-cellular junctions than bi-cellular junctions, as seen in Fig. 4b. This shows that particles with larger size transmigrate through tri-cellular junctions and those with smaller size transmigrate via bi-cellular junctions. We also study the effect of the varying magnitude of shear between cells and find that as the magnitude increases, permeability also increases, for a given gap size threshold, as seen in Fig. 4c. These observations imply that the chance of transcellular migration occurring at tri-cellular junctions or bi-cellular junctions depends on the mechanical stimuli. When the physiology and functioning of the cells are such that they undergo shear between each other, permeability at a tri-cellular junction is higher than at a bi-cellular junction.

### 3.2 Permeability increases with Calcium concentration

As discussed earlier, cytoskeletal contractility directly affects VE-cadherin bonds. It was seen that Rho/ROCK activation plays a critical role in regulating the endothelial barrier function ^11^. Since our mathematical model considers the effect of actin-contractility via calcium concentration *C*^^^, we study the effect of contractility on permeability by varying *C*^^^. Upon its increase, stress fibre concentration *η* increases leading to an increased active stress *σ_a_*. This results in cell-cell contact experiencing higher traction that leads to loss of contact and results in higher permeability, as can be seen in Fig. 5. Low levels of *C*^^^ indicate that the contractile stress is not high, and permeability is dominated by the random forces acting on the contact surfaces. It can be seen that for *C*^^^ = 1, permeability is around 0.55. This implies that 55% of the junctions are permeable for a particle of size less than or equal to 0.15*µ*m. By changing the parameter values of cell-cell junctions and active stress growth, low and high levels of permeability a cell monolayer exhibits can be changed, as and when required to simulate specific experimental observations. Growth of permeability over time for different values of *C*^^^ is given in supplemental material section S8, Figure S5.

**Figure 5:**
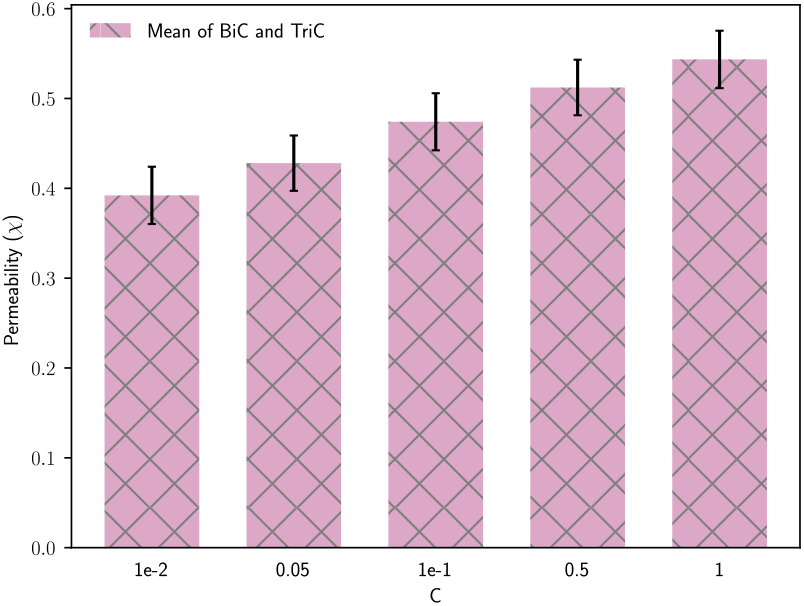
Effect of *C*^^^ on permeability. With an increase in *C*^^^, higher active stress leads to increased permeability.

### 3.3 Permeability increases with ECM stiffness due to mechano-chemical coupling

Recent experimental studies indicate that cardiovascular and many other diseases are associated with variations in the mechanical properties of the ECM. Biomechanical tests have found that atherosclerotic plaques are stiffer than endothelial basement membranes by several orders of magnitude ^40, 41^. Mechanical properties of ECM are also found to be responsible for Endothelial-Mesenchymal transition (EndMT), which is a characteristic of cancer extravasation occurring via endothelium exhibiting abnormal permeability ^42^. In order to study permeability, experiments *in-vitro* involve culturing cells on a substrate and using pro-inflammatory agents to induce permeability. Several *in-vitro* experiments have shown the relation between substrate properties, both mechanical ^14, 43^ and geometrical ^15, 44^ on permeability. Cells, when attached to the ECM, form focal adhesions consisting of integrins, which transfer forces to and from the cell. Downstream, focal adhesions are connected to the cytoskeleton machinery, which is further connected to VE-cadherins. Hence, VE-cadherin behaviour is regulated by integrin behaviour and vice-versa. Accordingly, it has been observed that with an increase in ECM stiffness, the contractility of the cells increases ^45^, and also the permeability of the monolayer ^14^.

In this regard, we consider an ECM below the planar monolayer, as shown in Fig. 6. For simplicity, in this article, the ECM is considered to be linear elastic. To begin with, we change the ECM stiffness and keep all other parameters fixed to find that permeability reduces with an increase in stiffness, as shown in Fig. 7a. This reduction is due to the higher equivalent stiffness of cell-ECM bond with higher ECM stiffness. But, experimental observations showed that permeability increases with the stiffness of ECM ^14^. Hence a multi-parameter analysis was carried out by changing *C*^^^ and ECM stiffness simultaneously, as shown in the heatmap Fig. 7a. It could be seen that neither a purely mechanical effect (row-wise comparison) nor a purely chemical effect (column-wise comparison) simulate the right behaviour of cells. It is to be noted that here even though a purely chemical effect exhibits increasing permeability, it would be independent of ECM stiffness, which we consider here as a mechanical effect. Hence a mechano-chemical coupled model is necessary to simulate the behaviour of cells to varying ECM stiffness (diagonal elements in the heatmap). In this regard, we can conclude that with an increase in ECM stiffness, cells respond by increasing active stress via higher contractility leading to higher permeability, as shown in Fig. 7b.

**Figure 6:**
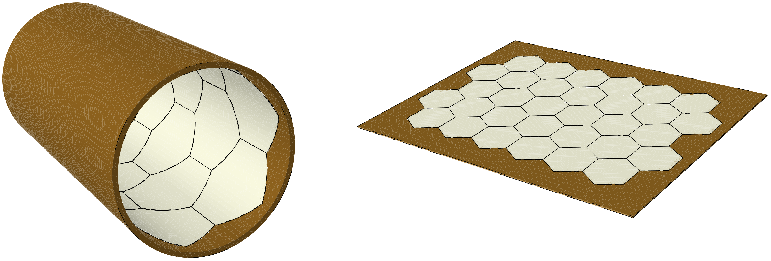
Planar and cylindrical endothelial monolayers attached to the substrate.

**Figure 7:**
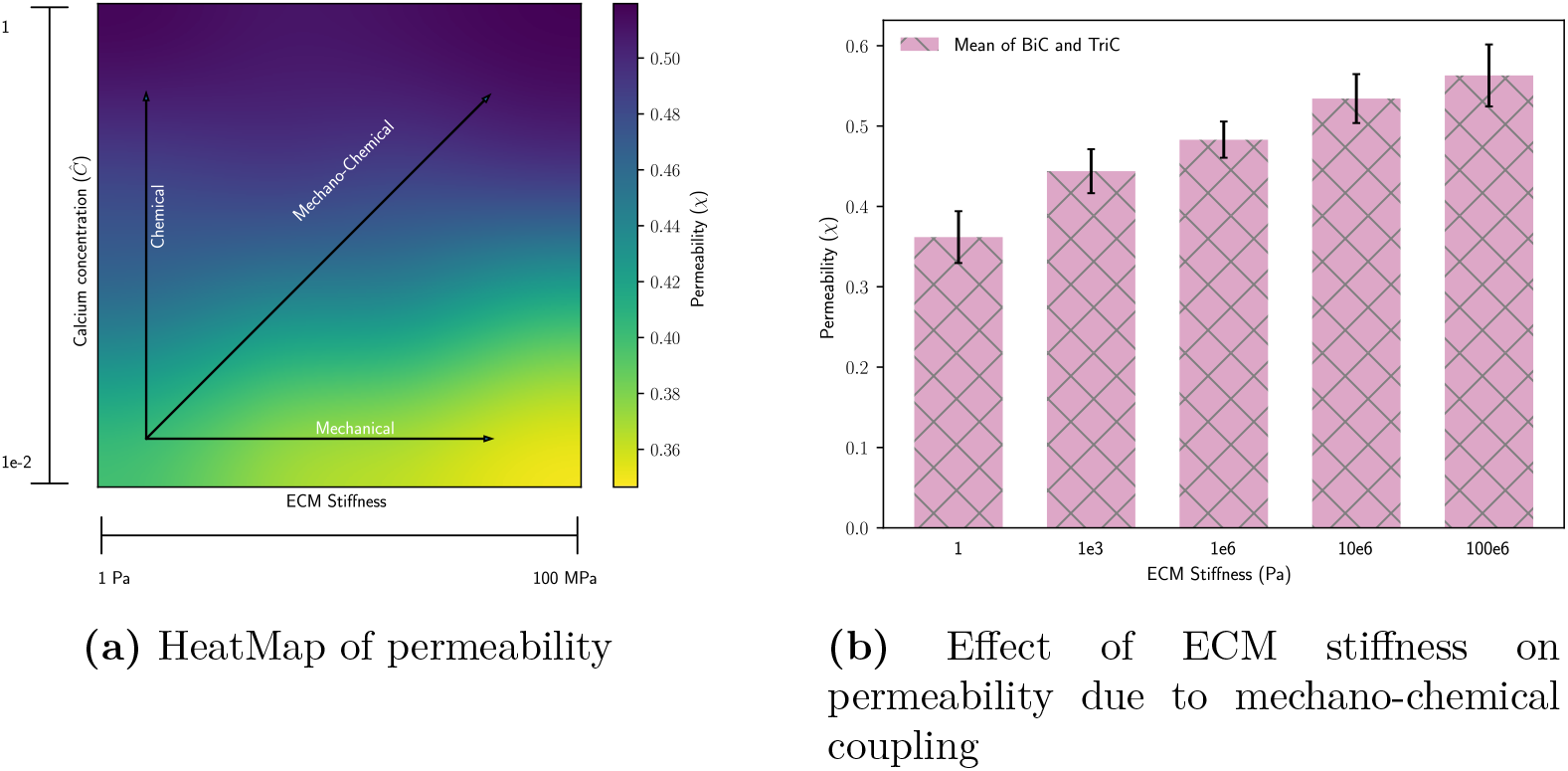
Effect of ECM stiffness on permeability is a mechano-chemical effect. a) Heatmap of permeability for purely mechanical(rows), purely chemical(columns) and mechano-chemical (diagonal) effect. An increase in ECM stiffness along with an increase in calcium concentration which induces higher contractility is necessary to simulate the effect of ECM stiffness on permeability. b) With an increase in ECM stiffness along with mechano-chemical coupling (diagonal elements), the monolayer exhibits higher permeability.

### 3.4 Traction due to blood flow increases permeability with ECM stiffness

Micro vessels *in-vivo* experience several stimuli simultaneously; shear stress, hemodynamic pressure, the tension on the walls and other local stresses due to geometric effects. In a healthy vascular network, the mechanical equilibrium of endothelial cells, under the influence of such forces, is necessary for its proper functioning. Any changes to the equilibrium lead to physiological growth and remodelling in order to maintain homeostasis ^46^. The inability to remodel the tissues results in an undesired stress state and makes individuals susceptible to cardiovascular diseases (CVDs) ^47^. Atherosclerosis is one of the very common CVDs and is responsible for the death of millions of people across the world. Abnormal permeability in the endothelium results in the deposition of fat molecules, and cholesterol in between the endothelium and surrounding smooth muscle cells ^48^. This results in the formation of atherosclerotic plaques that are very stiff and reduces the diameter of the vessel causing obstruction to the flow of blood in the blood vessel. It has been observed in several *in-vivo* and *in-vitro* experiments that atherosclerotic plaques are mainly found in the region where blood flow is disturbed, which is common in branching points of the vessel ^49, 50^. Thus, we study the effect of random traction forces on increasing substrate stiffness, simulating the effect of disturbed flow in the regions where atherosclerotic plaques are found. We find that with an increase in the stiffness of the substrate and the magnitude of traction forces that cells experience, permeability increases as well. For a constant ECM stiffness, an increase in traction magnitude increases the permeability. But this effect is found to be enhanced when the substrate stiffness is higher, as shown in the heatmap Fig. 8. This suggests that as the traction forces increase upon reduction in the diameter of the blood vessel, permeability increases leading to an increased plaque formation. The higher stiffness of plaques further increases permeability, thereby causing a positive feedback loop.

**Figure 8:**
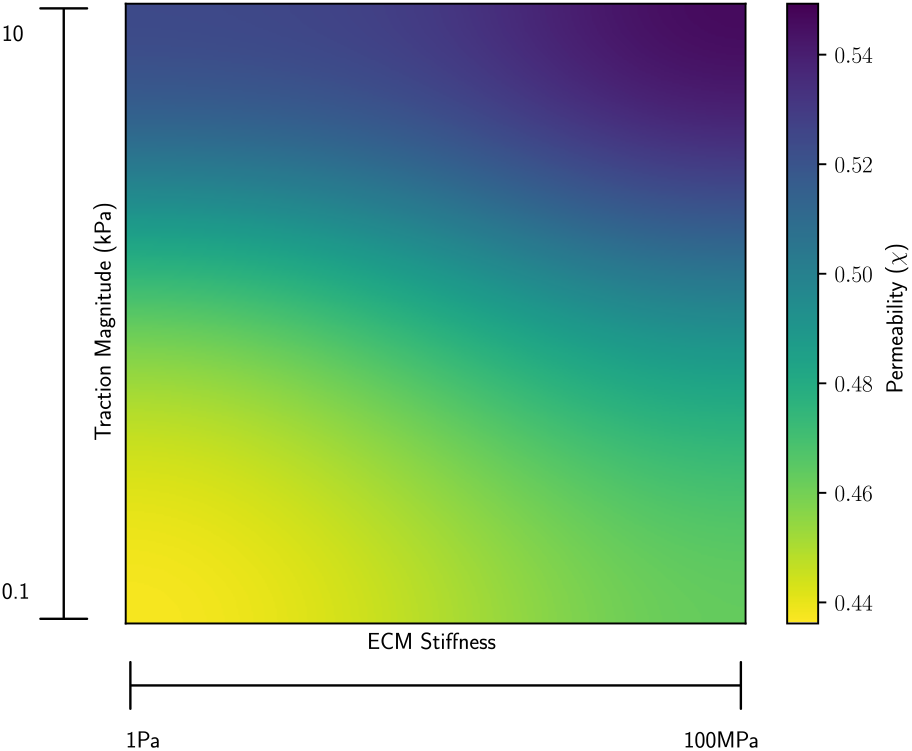
Permeability increases with increasing traction forces and substrate stiffness, indicating the positive feedback loop between permeability and ECM stiffness.

### 3.5 Cylindrical monolayer exhibits higher permeability than planar monolayer

Usually *in-vitro* experiments are performed by seeding endothelial cells on a petri-dish or a glass slide where they form a planar monolayer. But *in-vivo*, endothelial cells form cylindrical-shaped vascular tubes that allow for the transport of blood. In addition, they are surrounded by ECM that alters the properties and behaviour of endothelium. Hence, in order to extrapolate experimental observations from a planar monolayer to a vascular tube, that is in order to be able to predict *in-vivo* behaviour based on *in-vitro* experiments, we need to understand the difference in the behaviour of cells due to changing environmental conditions. Using the model presented in this article, we can simulate the behaviour of endothelial cells forming a micro-vessel and study the effect of ECM and other multi-physics loading on endothelial permeability.

To begin with, we form a cylindrical monolayer with 3D endothelial cells as shown in Fig. 6, but without any ECM surrounding it. The ends of the vessel are fixed to prevent rigid body motion. By only applying random pressure loading between cells as described earlier, we can measure permeability at bi-cellular and tri-cellular junctions. Simulations show that cylindrical monolayer exhibits higher permeability than planar monolayer for the same set of parameters, as can be seen in Fig. 9. Compared to planar monolayer, cylindrical monolayer took longer to reach the steady state, and the value was higher as well. We observed that higher permeability arises mainly due to the additional degree of freedom that cells enjoy due to curved geometry.

**Figure 9:**
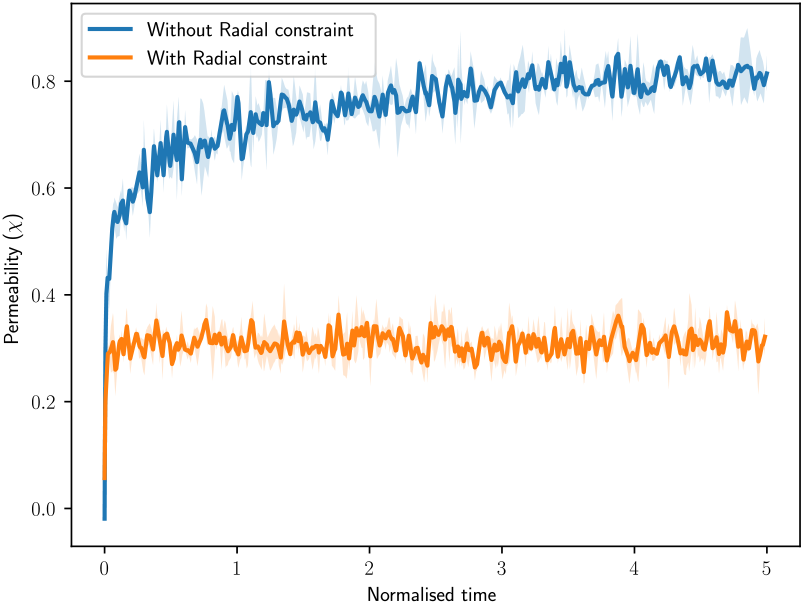
Permeability in a cylindrical geometry with and without radial constraint. It is found that cylindrical monolayer without any ECM constraint exhibits higher permeability than the planar monolayer (Fig. 3a)

By restraining this degree of freedom, which is equivalent to a rigid ECM surrounding the cylindrical monolayer, we see that the permeability of the cylindrical monolayer is similar to that of a planar monolayer Fig. 9. These simulations show that permeability observed in *in-vitro* experiments performed on a planar monolayer of endothelial cells is not directly applicable to *in-vivo* conditions. The results from this model are purely qualitative in nature and need to be quantified against experimental observations performed on cylindrical monolayers, and the effect of ECM properties could be studied as well.

## 4 Conclusion, and future work

In this article, we have introduced a novel continuum-level modelling framework for studying the dynamic behaviour of adherens junctions regulated by VE-cadherins. Traction-separation law as a phenomenologically equivalent mechanism for catch-slip bond law has been used to model the opening and closing of cell-cell junctions. The stiffness of the VE-cadherin bond, the stretch upon which the VE-cadherin bond dissociation is initiated and completely dissociated are the parameters needed to define the law, which are directly available in the literature ^30^. The cytoskeleton of endothelial cells is assumed to be made up of linear elastic passive component and non-linear strain-rate dependent active component. The ODE for stress fibre concentration growth determines the level of active stress in the cytoskeleton. Through the mechanical equilibrium between the stress in the cytoskeleton and the force applied by cell-cell contact and cell-extracellular matrix (ECM), mechanical feedback between them can be established.

We define permeability as the ratio of the number of open junctions to the total number of adherens junctions present in the monolayer. The model has been able to predict the variation in permeability due to varying gap sizes and multiple mechanical stimuli acting simultaneously. Numerical simulations can be performed to study how using dyes of different sizes show varying levels of permeability. The size of the gap present in the monolayer might be smaller than the molecular size of the dye resulting in reduced extravasation. Thus, the model can be used to correlate the quantification of VE-cadherin staining and permeability. Shear stress between the cells is found to play a huge role in VE-cadherin bonds and hence the permeability of the monolayer. Since shear stress varies with endothelial cell types, the simulations give an insight into differences observed in *in-vitro* experiments performed with different cell types and passage numbers. Thus, through the *in-silico* experiments, we are able to develop testable hypotheses on the conditions that result in higher permeability. Simulations also showed that permeability increases with calcium concentration-induced contractility. The role of substrate stiffness on permeability was studied by attaching cells to a linear elastic substrate. It was seen that an increase in substrate stiffness results in lowering the permeability. But a simple mechano-chemical coupling was necessary to show that permeability increases with substrate stiffness. Even though the mechano-chemical coupling used in this article is a very simple one, where *C*^^^ is varied linearly with ECM stiffness, this provides a perfect platform to further study the role of varying mechanical and geometric properties of the substrate on permeability. The linear elastic substrate could be further replaced by collagen-type fibrous material and a detailed mechano-chemical coupling pathway could be added easily.

It was interesting to note that the geometry of blood vessels determined the probability of the location of atherosclerotic plaques. They were commonly found near the branches of valves where the blood flow is in a disturbed state ^51^. In numerical simulations, we could study the effect of such a disturbed flow on permeability and its dependence on ECM stiffness by applying random traction forces on endothelial cells in contact with a substrate. Results showed that applying random traction forces increases permeability, and this effect was higher with increasing ECM stiffness.

Studies also showed that the cylindrical monolayer, which is similar to endothelium *in-vivo*, exhibits higher permeability than the planar monolayer, when there is no ECM surrounding it and behaves like a planar monolayer when surrounded by a rigid ECM. Thus, the hypothesis that we develop from the simulations is that the experiments performed *in-vitro* on a planar monolayer do not directly correlate to the behaviour *in-vivo*, and the geometry of the endothelium plays a prominent role in determining permeability.

The model could be thus used to understand the role of chemical and mechanical cues, and test the conditions resulting in extravasation of leukocyte *in-vivo* or a dye *in-vitro*. This can also be used to test under what conditions drug molecules can extravasate, and through either tri-cellular or bi-cellular junctions. This can have several important applications in drug delivery. The model could also be used to determine the right mechanical properties of ECM that can regulate permeability. This could be used as a method to stop the positive feedback between ECM and cell contractility resulting in reduced permeability. While the current treatment techniques involve changing the properties of blood using drugs such as statins or using stents to regulate the diameter of arteries ^48^, the model could be particularly used in finding the right mechanical and chemical environment necessary for either maintaining dynamic homeostasis or rectifying the properties of ECM such that the cells go back to a healthy state. In addition to atherosclerosis, this model could be used to bolster the antifibrotic treatment techniques used in several diseases such as cancer, and liver diseases ^52^, which helps in regulating permeability of endothelium and thereby maintain vascular homeostasis. Further, the model could be used to effectively engineer vascular network *in-vitro* by studying the role of hydrogel on vascular permeability ^53^. The model could also in future be extended to understand the role of ageing on vascular stiffening ^54^ or the effect of cytoskeletal metabolic memory ^55^ on cell-cell junctions and thereby permeability. The phenomenological model presented in this article is motivated by the mechanical responses observed during *in-vitro* experiments. The formulation of the model is such that it is possible to extend it to include detailed mechano-chemical coupling pathways that affect permeability. The traction-separation law which has displacement as the only factor determining when the bonds dissociate completely can include the concentration of VE-cadherin or other proteins as the additional factor that determines the dissociation of VE-cadherin bonds. Simulations showed that permeability was slightly higher in the regions experiencing disturbed flow-type traction forces compared to regions where traction forces are uniform, shown in the supplemental material section 9, Figure S6. We believe that the low increase in permeability with disturbed flow-type conditions is due to the absence of mechanosensors such as glycocalyx that further regulate the cytoskeleton downstream, and vary permeability ^56, 57^. Thus, the model presented in this article provides a perfect platform for several extensions and mechano-chemical studies.

## Acknowledgements

We would like to acknowledge the funding from BBSRC grant [BB/V002708/1], and UKRI Future Leaders Fellowship [MR/T043571/1]. We would also like to thank Giovanni Guglielmi for discussions about using friendly colours in plots.

