## Supplemental for "A mechanical modelling framework to study endothelial permeability"

### S1 Source files

Link to download all the source files:
https://github.com/bkprdp/VE-Cadherin-Mechanical-Model-Abaqus
File descriptions :

1. Cell_UMAT_Small_Strain.f: User-defined material. Active and Passive stress growth, using small strain formulation.

2. UAMP : Random pressure amplitude

3. Planar_Monolayer_Without_ECM.inp : inp file for planar monolayer without ECM

4. Planar_Monolayer_With_ECM.inp : inp file for planar monolayer with ECM

5. Cylindrical_Monolayer_Without_ECM.inp : inp file for cylindrical monolayer without ECM

6. Cylindrical_Monolayer_With_Rigid_ECM.inp : inp file for cylindrical monolayer with fixed radial constraint representing rigid ECM

7. Nodes_Bi_Tri_Cellular_Jncs.py : Python script to classify nodes belonging to bi-cellular and tri-cellular junctions

8. Planar_Monolayer_COPEN_Analysis.py : Python script to perform quantitative analysis of the csv file containing copen values

Link to view animations of simulations:
https://drive.google.com/drive/folders/1qoBRa-cLU5RXyTTd6mylStY4yN5dRakC?usp=drive_link

### S2 Traction-Separation law

In this article, traction-separation law is hypothesised to be the mechanical equivalent of the catch-slip bond law. The parameters of traction-separation law could be varied to modify the association and dissociation behaviour of VE-cadherin bonds. As explained in the main text, only three parameters are needed :

1. Stiffness of VE-cadherin bond

2. Stretch at which the VE-cadherin bond starts to dissociate

3. Stretch at which the VE-cadherin bond is completely dissociated

It can be seen in Figure S1, as the normalised stiffness is increased from 1 to 2, the maximum contact force that the bond can take is also doubled. When the maximum separation that the bond can handle before complete dissociation is increased, the time the bond takes to dissociate also increases. It can be easily seen that by changing $\delta_i^0$, stiffness can be changed as well. Thus, by varying these parameters, quantitative comparison with experiments is possible.

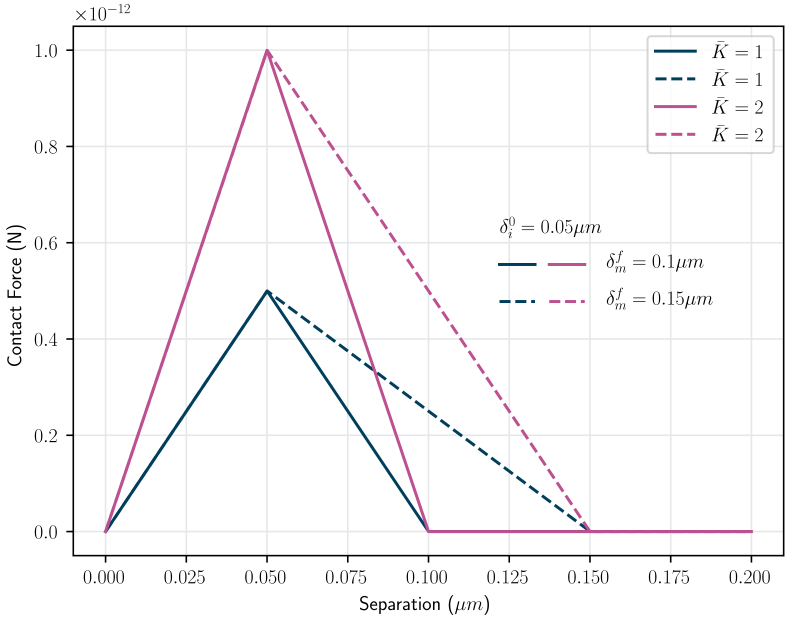


**Figure S1:** Parameters involved in traction separation law can be varied to study the maximum force, time of association and dissociation of VE-cadherin bonds.

### S3 Cauchy stress tensor derivation

Any unit vector $\mathbf{m}$ along the fibre direction ($\omega$,$\phi$), Fig. S2, can be written as

$$\mathbf{m} = sin(\omega)cos(\phi)\mathbf{x_1} + sin(\omega)sin(\phi)\mathbf{x_2} + cos(\omega)\mathbf{x_3} \tag{1}$$

Following [1], we can write the components of active Cauchy stress tensor as

$$\sigma_{ij}^a = \frac{3}{4\pi}\int_0^{2\pi}\int_0^{\pi} \sigma^a(\omega,\phi)m_i m_j sin(\omega)d\omega d\phi \tag{2}$$

where i,j = 1,2,3. In the case of 2D geometry,

$$\sigma_{ij}^{a} = \frac{1}{\pi} \int_{-\pi/2}^{\pi/2} \sigma^{a}(\phi) m_i m_j d\phi \tag{3}$$

where i,j=1,2.

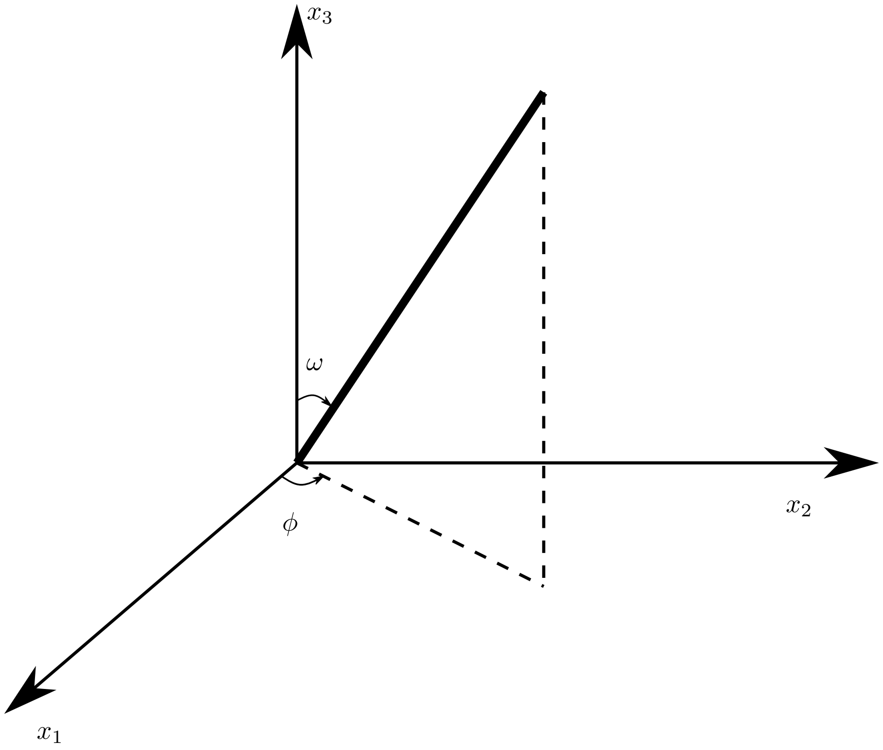


**Figure S2:** Orientation of fibre with respect to $x_1$, $x_2$, and $x_3$ axes.

### S4 Finite Strain formulation

In this article, all the simulations presented used small strain formulation. This can be further extended to finite strain formulation using non-linear hyper-elastic materials. Here an example with Neo-Hookean material is given. Passive stress can thus be formulated with the strain energy function

$$\psi(I_1, J) = C_{10}(I_1 - 3) + \frac{1}{D_1}(J - 1)^2 \tag{4}$$

where $\psi$ is the strain energy density function, $I_1$ is the first invariant of the left Cauchy-Green tensor $\mathbf{B}$, $J$ is the determinant of the deformation gradient

$\mathbf{F}$. $C_{10}$ and $D_1$ are the material parameters related to Young's modulus and Poisson ratio via Bulk modulus ($K_0$) and shear modulus ($\mu_0$) as given in Eq. (5).

$$\begin{aligned} C_{10} &= \frac{\mu_0}{2} \\ D_1 &= \frac{2}{K_0} \end{aligned} \tag{5}$$

Left Cauchy-Green tensor is defined as

$$\mathbf{B} = \mathbf{F}.\mathbf{F}^{\mathbf{T}} \tag{6}$$

The constitutive equation is given by the first derivative of the strain energy function

$$\sigma_{ij}^{p} = \frac{2}{J} C_{10} \left( \bar{B}_{ij} - \frac{1}{3} \delta_{ij} \bar{B}_{kk} \right) + \frac{2}{D_1} (J-1) \delta_{ij} \tag{7}$$

where $\delta_{ij}$ is the Kronecker delta, and $i, j = 1, 2, 3$. $\bar{\mathbf{B}}$ is the volumetric strain tensor given by

$$\bar{\mathbf{B}} = J^{\left(\frac{-2}{3}\right)} \mathbf{B} \tag{8}$$

In general, the Jacobian matrix $\mathbf{C}$ needed for the Newton scheme used by ABAQUS, can be obtained from the equilibrium equation

$$\delta \boldsymbol{\tau} = J \mathbf{C} : \delta \mathbf{D} \tag{9}$$

where $\boldsymbol{\tau}$ is the Kirchoff stress, related to the Cauchy stress as $J\boldsymbol{\sigma}$. The rate of deformation $\mathbf{D}$ is defined as

$$\delta \mathbf{D} = \mathbf{sym} \left( \delta \mathbf{F}.\mathbf{F}^{-\mathbf{1}} \right) \tag{10}$$

Solution of Eq. 9 can be obtained numerically by perturbing the individual component of deformation gradient $\mathbf{F}$ by a small amount $\epsilon$ as described in [2, 3]. Derivation leads to

$$\mathbf{C} = \frac{1}{J\varepsilon} \left[ \boldsymbol{\tau} \left( \hat{\mathbf{F}} \right) - \boldsymbol{\tau} \left( \mathbf{F} \right) \right] \tag{11}$$

where $\hat{\mathbf{F}}$ is the perturbed deformation gradient given as

$$\hat{F}_{ij} = F_{ij} + \Delta F_{ij} \tag{12}$$

In addition, in the case of finite strain formulation, the rotation of fibres have to be taken into account during the evaluation of active stress tensor. Thus (2) becomes

$$\sigma_{ij}^{a} = \frac{3}{4\pi} \int_0^{2\pi} \int_0^{\pi} \sigma^a(\omega, \phi) m_i^* m_j^* sin(\omega) d\omega d\phi \tag{13}$$

where $m_i^*, m_j^*$ are the vectors in the current configuration evaluated as

$$m_i^* = \mathbf{R} m_i \tag{14}$$

where $\mathbf{R}$ is the rotation matrix evaluated via the decomposition of the deformation gradient as

$$\mathbf{F} = \mathbf{R}\mathbf{U} \tag{15}$$

### S5 Parameter values

Parameter values used in this article are given in Tab. S1. Some of the parameter values are obtained directly from the literature while some of them are modified to obtain qualitatively comparable results with the experiments.

| Parameters | Value | Units | Reference |
|---|---|---|---|
| Passive stiffness | 10 | kPa | [4] |
| Poisson ratio | 0.45 | - | - |
| Max stress in stress fibre | 1 | MPa | [5] |
| Strain rate coefficient | 0.1 | /s | [5] |
| $k_f$ | 0.01 | /s | [5] |
| $k_b$ | 0.1 | /s | [5] |
| Bond stiffness | 1 | N/m | [6] |
| Damage initiation | 0.05 | $\mu$m | [6] |
| Damage termination | 0.15 | $\mu$m | [6] |
| Cadherin Concentration | 1e7 | cadherins/m$^2$ | - |

**Table S1:** Baseline parameter values

### S6 Convergence study

Mesh convergence analysis is performed. It can be seen in Fig. S3 that the error (evaluated wrt to results from a very fine mesh simulation) with a coarse mesh is 5%. We assume that this error is acceptable for the qualitative nature of this article. Hence all the analyses performed in this article were performed with the mesh with an element size of 1$\mu$m.

### S7 Uniform vs Random gap opening

A comparison of gap opening, when the contraction is uniform and random, is shown in Fig. S4. The simulated animations of the deformation of the endothelial monolayer, in case of uniform and random loading, can be found in the link provided in supplemental section S1

### S8 Effect of $\hat{C}$ on permeability

As Calcium concentration is increased, contractility increases resulting in higher permeability. Variation of permeability over simulation time is given in Figure S5.

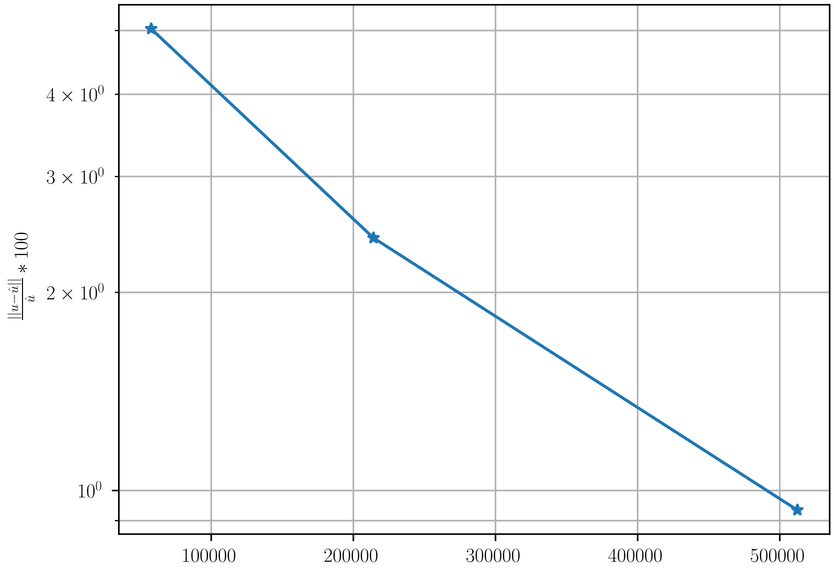


**Figure S3:** Mesh convergence study

### S9 Disturbed flow vs uniform flow

We define permeability as the ratio of the number of open junctions to the total number of junctions in the monolayer. As expected, when the shear stress gradient between cells increases, the probability of cell-cell junctions being open increases. In this regard, we saw that in disturbed flow conditions, where the flow is randomly varying in both x and y directions, permeability was slightly higher than that of a uniform flow condition, where all cells experience the same flow, as shown in Fig. S6. It is to be noted that the current analysis shows that the difference between uniform and disturbed flow is very small. This might be either due to the values of parameters chosen in this study or due to the lack of specific mechano-sensory channels that can sense flow such as glycocalyx. Thus, in future, this analysis will be coupled with glycocalyx which is found to play an important role in regulating permeability as well [7]. Further development could help us in understanding the role of flow in diseases such as atherosclerosis and other permeability-related diseases.

#### S10 Thickness of cell

In the article, we consider the thickness of the cell as $0.1\mu$m. But in literature, it has been found that thickness varies till 10 $\mu$m. In this regard, we varied the thickness of the cell and studied its effect on permeability. As the thickness of the cell is increased, keeping the cadherin concentration constant, the net force that cell-cell contact can resist without damage increases. But simultaneously, the contractile force and force due to random pressure loading on the boundary also increase. Hence the contributions balance out and we do not observe any difference in permeability due to changing thickness of the cell as can be seen in Fig. S7. In future, this study could be extended by considering the effect of the nucleus, which will alter cell stiffness locally.

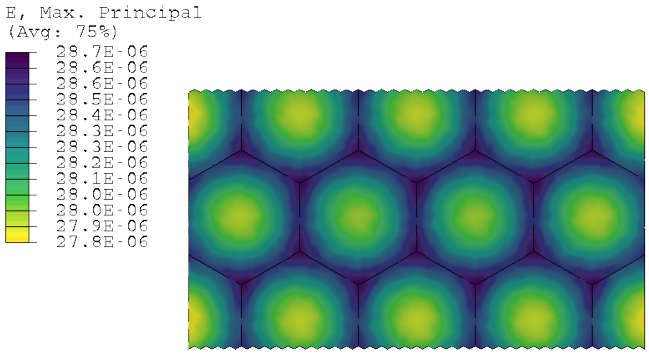


**(a)** Strain distribution due to uniform contraction of cells.

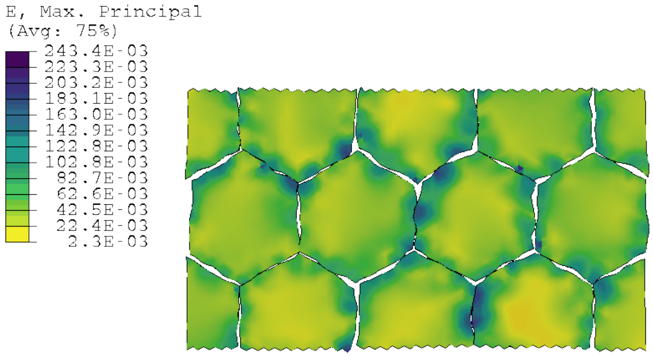


**(b)** Strain distribution due to random pressure loading.

**Figure S4:** Strain field in a planar monolayer subjected to different types of loading. a) Due to the uniform contraction of cells, strain is high at a tri-cellular junction compared to bi-cellular junction. b) Due to random loads between cells, strain is high either at a bi-cellular or tri-cellular junction depending on the load.

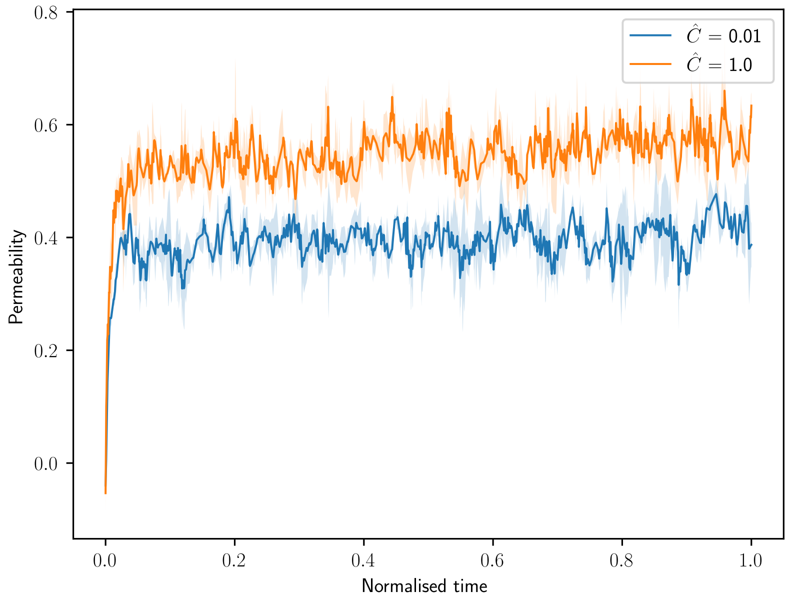


**Figure S5:** Variation of permeability over time

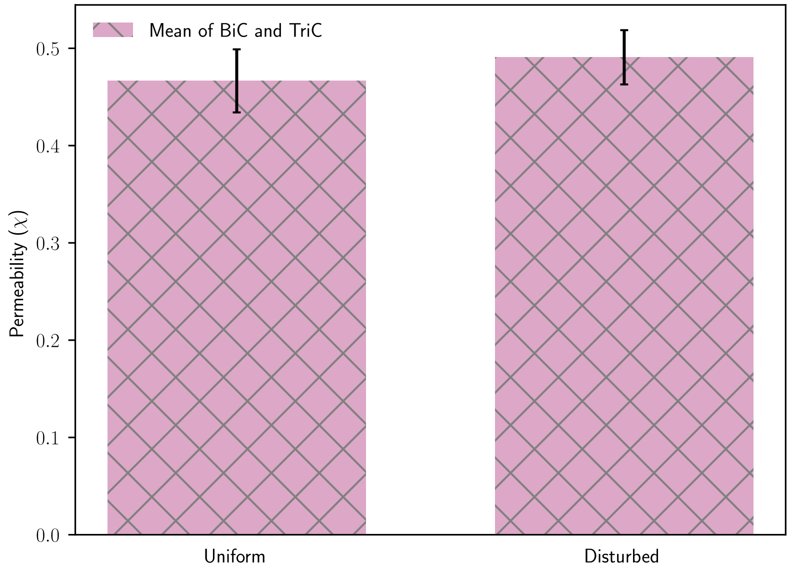


**Figure S6:** Variation in permeability due to uniform and random flows

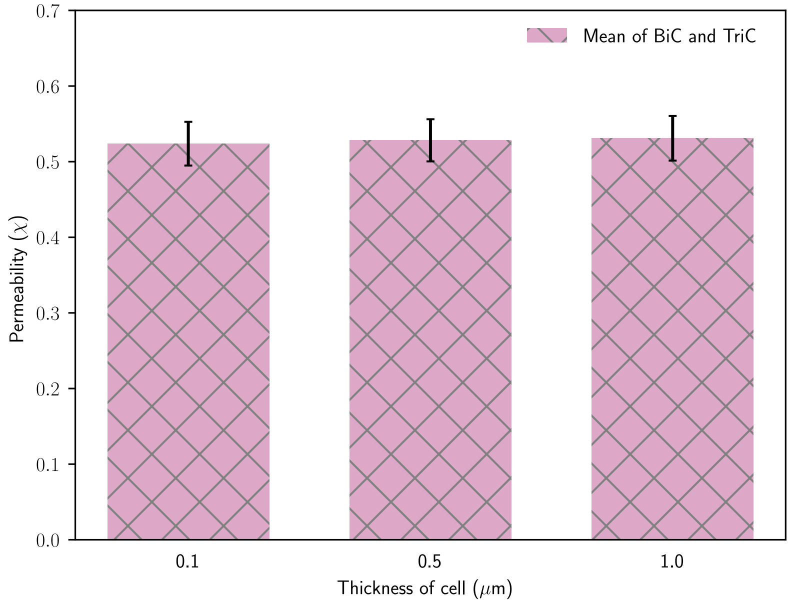


**Figure S7:** Variation in permeability due to change in thickness of cell
